# Formation and maintenance of robust long-term information storage in the presence of synaptic turnover

**DOI:** 10.1101/023531

**Authors:** Michael Fauth, Florentin Wörgötter, Christian Tetzlaff

## Abstract

A long-standing problem is how memories can be stored for very long times despite the volatility of the underlying neural substrate, most notably the high turnover of dendritic spines and synapses. To address this problem, here we are using a generic and simple probabilistic model for the creation and removal of synapses. We show that information can be stored for several months when utilizing the intrinsic dynamics of multi-synapse connections. In such systems, single synapses can still show high turnover, which enables fast learning of new information, but this will not perturb prior stored information (slow forgetting), which is represented by the compound state of the connections. The model matches the time course of recent experimental spine data during learning and memory in mice supporting the assumption of multi-synapse connections as the basis for long-term storage.

**Author Summary:** It is widely believed that information is stored in the connectivity, i.e. the synapses of neural networks. Yet, the morphological correlates of excitatory synapses, the dendritic spines, have been found to undergo a remarkable turnover on daily basis. This poses the question, how information can be retained on such a variable substrate.

In this study, using connections with multiple synapses, we show that connections which follow the experimentally measured bimodal distribution in the number of synapses can store information orders of magnitude longer than the lifetime of a single synapse. This is a consequence of the underlying bistable collective dynamic of multiple synapses: Single synapses can appear and disappear without disturbing the memory as a whole.

Furthermore, increasing or decreasing neural activity changes the distribution of the number of synapses of multi-synaptic connections such that only one of the peaks remains. This leads to a desirable property: information about these altered activities can be stored much faster than it is forgotten. Remarkably, the resulting model dynamics match recent experimental data investigating the long-term effect of learning on the dynamics of dendritic spines.

## Introduction

Learning and memory in neuronal networks is commonly attributed to changes in the connectivity, i.e. the number of synapses between neurons and their transmission efficacy [1, 2, 3], which are then consolidated to form long lasting memories [4, 2, 5, 6]. The majority of the excitatory cortical synapses resides on dendritic spines [7]. Thus, very likely they contribute much to these phenomena.

One central problem here is the apparent contradiction between the facts that memories are often stable over very long times, while the underlying neural substrate - most notably the synaptic spine structure - undergoes constant changes.

Experiments reveal that even under control conditions, more than 5% of the spines are exchanged every day [8, 9, 10, 11, 12, 13]. This turnover increases during learning (e.g, motor learning [11, 12]), sensory deprivation [13] or environmental enrichment [12].

Intriguingly, although learning triggers increased turnover, *repeated* learning of the same task – even after long intervals – does not. The missing need for restructuring during the repeated training has been attributed to structural changes that had persisted from the previous training, and this, in turn, has been interpreted as a long-lasting memory trace [11, 12, 13]. This trace can be acquired during a few days of training, but maintained for several months, which indicates different time scales for learning and forgetting. However, the majority of the spines formed during learning vanishes a few days after training and only very small proportions may persist for the survival duration of the memory trace. Thus, it seems more reasonable that the collective properties of the pool of synapses, rather then any momentarily existing connectivity configuration, have to account for learning and memory formation.

Several such properties have been found, such as the abundance of two- or three-neuron microcircuits [14, 15] or the characteristic distribution of the number of synapses between two neurons, which shows a large peak at zero synapses and another smaller peak at multiple synapses [16, 17, 18, 19, 20]. Theoretical results suggest that such bimodal distributions emerge from the collective dynamics of multiple synapses [21].

One possibility to obtain this collective dynamics is to establish a positive feedback between the number of synapses and synapse stability. For this, however, each synapse has to sense the number of synapses from the same presynaptic neuron. It has been shown theoretically that the postsynaptic activity, which can be accessed by each synapse, provides sufficient information [22, 23]. However, only a complex non-linear interaction between structural and synaptic plasticity, i.e. between the number of synapses, the postsynaptic activity, the synaptic weights, and synapse stability, gives rise to the observed bimodal distributions [22, 23]. For this, synaptic plasticity rules must be sensitive to either correlations of pre- and postsynaptic firing [22] or to the postsynaptic firing rates [23].

A systematic variation of the neural activity in such systems reveals that the bimodal distribution emerges from a hysteresis in the creation and deletion of synapses [23]. The existence of such a hysteresis could be used to store information [24, 25, 26, 27, 28, 29, 23].

Considering these results, here, we abstract this collective dynamics by using a theoretical model in which the stability of synapses directlydepends on the actual number of synapses. We demonstrate that storing information in the collective dynamics of multiple synapses can resolve the contradiction between memory duration and structural/synaptic volatility. Furthermore, we show that especially bimodal synapse distributions correspond to a bistability in the number of synapses which enables prolonged information storage. These bistable systems allow fluctuations of the actual number of synapses within certain limits which keeps the structure flexible and leads to the intriguing additional effect that learning is orders of magnitude faster than forgetting. Recent experimental data on the dynamics of spines [11] can, therefore, naturally be explained by this model.

## Results

In neural systems information storage is commonly attributed to the connectivity based on synapses. Experiments reveal that a connection between two neurons can consist of multiple synapses – in the following named *compound-connection.* In this study we define a compound-connection between two neurons by two numbers: the number of potential synapses *N*, which accounts for the morphologies for the two connected neurons [30, 31, 23], and the number of realized synapses *S*, which are those potential synapses that actually host a functional contact. To mimic the turnover dynamics of dendritic spines from experiments, synapses are created and removed stochastically: New synapses are formed with a constant probability rate *b* at each of the potential (but not yet realized) synapses. Realized synapses are removed with a probability rate *d^ℑ^*[*S*], which depends on the number of realized synapses *S* and the current stimulation condition ℑ (Fig. 1 A).

**Figure 1.**
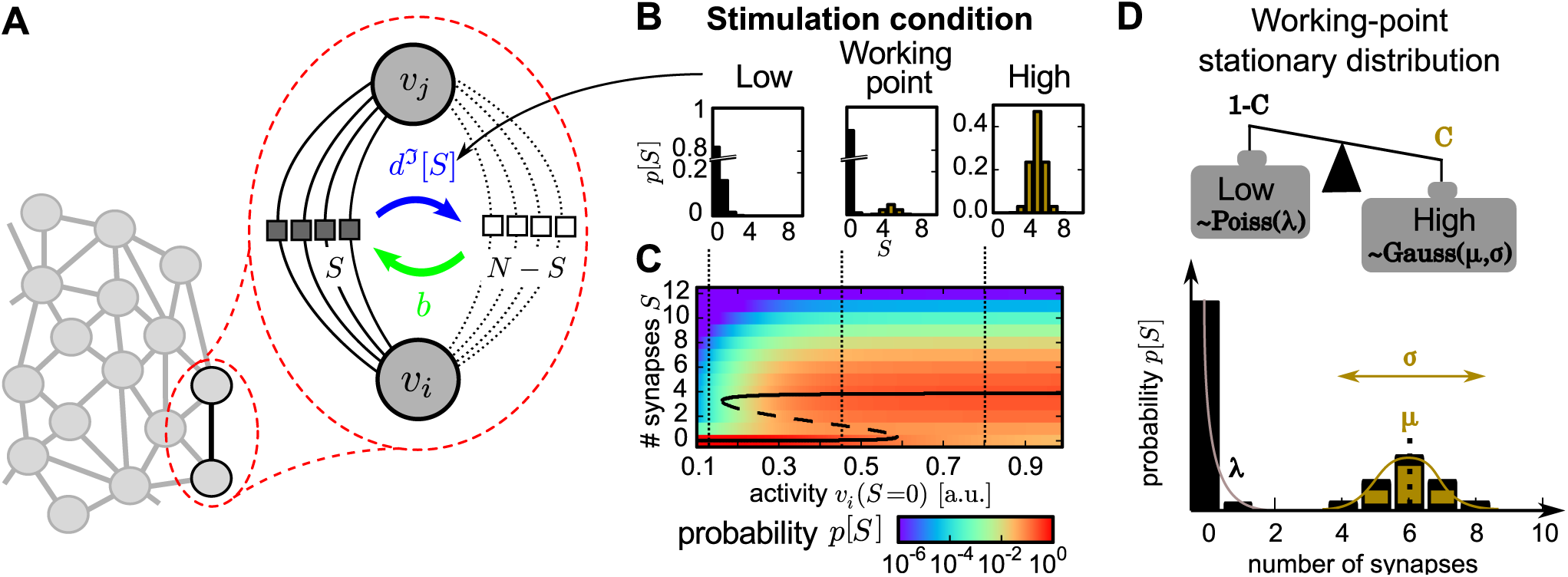
Model structure and the stationary distributions for different conditions. (A) Structure of the model: Each compound-connection consists of a number of potential synapses N and a number of realized synapses *S ≤ N*. Each realized synapse can be deleted with probability rate *d^ℑ^*[*S*] and, as long as *S ≠ N*, new synapses can be formed with constant probability rate *b*. (B) The dynamics of the model reveals three characteristic stationary distributions of synapses *p*[*S*] given different stimulation conditions *ℑ ∈* {*low, working-point, high*} (*low*: left; *working point* (*wp*): middle; *high*: right). (C) The stationary distributions (color code) emerge from the interaction of structural and synaptic plasticity when varying the stimulation (Figure adapted from [23]). Black line: theoretical positions of distribution maxima (solid) and minima (dashed). (D) The *wp-*stationary distribution consists of a weighted sum of the *high-* and *low*-distributions with weighting factors *C* and 1 − *C*. The *high*-distribution (brown) is modeled by a Gaussian distribution with mean *μ* and standard deviation *σ*, the *low*-distribution (black) by a Poisson distribution with mean and variance λ. Influence of distribution parameters is sketched schematically.

The dynamics of these processes can be described by a Markov process [22, 23]. Previous work, investigating their interaction with synaptic plasticity, shows that the stationary distributions of the number of synapses strongly depends on the stimulation an the resulting activity of the pre- and postsynaptic neuron (Fig. 1 C, [23]). In the current study, we abstract this to three characteristic activities, the so called *stimulation conditions* ℑ, yielding three qualitatively different stationary synapse distributions *p^ℑ^*[*S*] (Fig. 1 B): *low*-stimulation induces a low probability to form synapses and, therefore, leads to a unimodal distribution with peak at zero synapses, *high*-stimulation forms compound-connections consisting of mostly multiple synapses, and medium level stimulation, here called the *working-point-condition* (Fig. 1 D), leads to the working-point-(*wp-*)distribution, which is bimodal such as those found in many experimental studies [16, 17, 18, 19, 20]. Related to these findings, this is assumed to be the basic synaptic distribution to which compound connections converge at resting (medium) activity. Thus, *low*-stimulation corresponds to an – on average – reduced neuronal activity (possibly via inhibition) and *high*-stimulation to an excitatory input (possibly via external stimulation).

However, here, instead of modelling neural activity explicitly, these stationary distributions are more easily obtained by adjusting the deletion probabilities *d^ℑ^*[*S*] to render the different stimulation cases (see Models and Methods). This implementation yields bistable dynamics for the number of synapses under *wp-*conditions but only one stable fixed point under *low-* and *high*-conditions (compare Fig. 1 C dashed black line).

Using these different conditions, in the following we consider the dynamics of 500 compound-connections in our network and analyze the resulting probability distribution of synapses *p*[*S*].

Note all time scales, we refer to, are dependent on the synapse-formation probability rate *b*. However, from matching the model to dendritic spine experiments (see below; Fig. 6), we will demonstrate that the time scales of the model indeed correspond to memory-relevant scales of 100 days and more.

### Basic synaptic dynamics in the working point

First we simulated two groups of 500 compound-connections with deletion probabilities which enforce the development into the *working-point*-distribution. Hence, this corresponds to medium activities at all connections. However, we initialized both groups differently: the synapse distribution of the first group is from the beginning in the *wp-*distribution (Fig. 2 A), while in the second group all compound-connections initially consist of seven realized synapses (peak at *S* = 7; Fig. 2 B).

**Figure 2.**
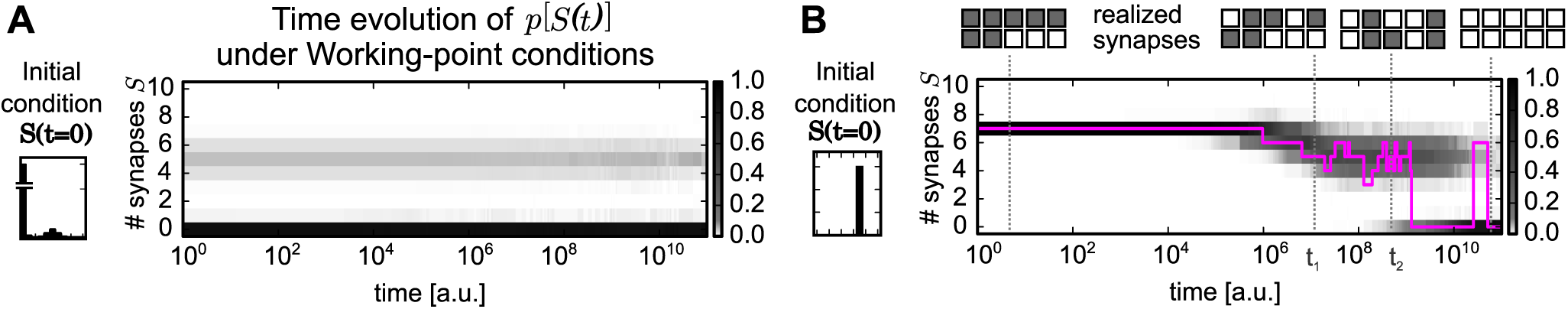
Temporal evolution of the probability distribution of synapses. The probability distribution *p*[*S*] of 500 compound-connections (gray scale) converges to the bimodal *wp-*stationary distribution given different initial conditions *p*[*S*(0)]: (A) initial distribution is the *wp-*distribution itself; distribution stay stationary (B) all compound-connections have seven realized synapses (*S* = 7); distribution converges to the *wp-*distribution. The development of a single compound-connection is shown as an example (purple).

The first group demonstrates the validity of the model setup as the system indeed remains stationary in the bimodal *wp-*distribution (Fig. 2 A).

The development of the second group (Fig. 2 B) reveals an interesting feature of the model dynamics: first, the compound-connections tend to delete synapses and, thus, the average number of synapses reduces from the initial value (*S* = 7) to the mean of the upper peak in the *wp-*distribution (*μ* = 5). In parallel the variance increases, thus, different compound-connections consist of different numbers of synapses. Later, although the probability distribution of the whole group of connections remains rather constant, the number of synapses of each single compound-connection varies frequently (see purple line for one example). Furthermore, even if during development the total number of realized synapses is the same at two different time points (e.g., *S*(*t*_1_) = *S*(*t*_2_) = 5), the location of these realized synapses could be different (Fig. 2 B, insets). However, the connection remains in the upper peak of the *wp-*distribution and fluctuates around the mean *μ*.

Finally, only after a long period (note the logarithmic time scale) many compound-connections delete all their synapses and a second peak at zero synapses emerges in the probability distribution. Even then – after reaching the *wp-*stationary distribution – the number of synapses at any given compound-connection fluctuates as can be seen, for instance, in the figure when the pink curve jumps between zero and six realized synapses. Such fluctuations within the stationary distribution (also in Fig. 2 A) represent the experimentally measured turnover of dendritic spines hosting a synapse [9].

Based on these results, we hypothesize that for long time scales the bistable dynamics of the number of synapses and the resulting probability distribution stores information rather than the existence of single synapses. This will be investigated next.

### Quantitative evaluation of information decay

In the second step, we investigate the capacity of the system to store information. For this, we evaluate how much information the number of synapses *S*(*t*) at time point *t* provides about the initial number of synapses *S*(0) using mutual information. Mutual information is an information theoretic measure, which determines how much of the variability of one quantity can be predicted by another one. Here, we calculate the mutual information between *S*(0) and *S*(t) (Fig. 3) for different initial probability distributions (Fig. 3 A1–3) and stimulation-conditions resulting in different stationary distributions (Fig. 3 B4, C4 and D4).

**Figure 3.**
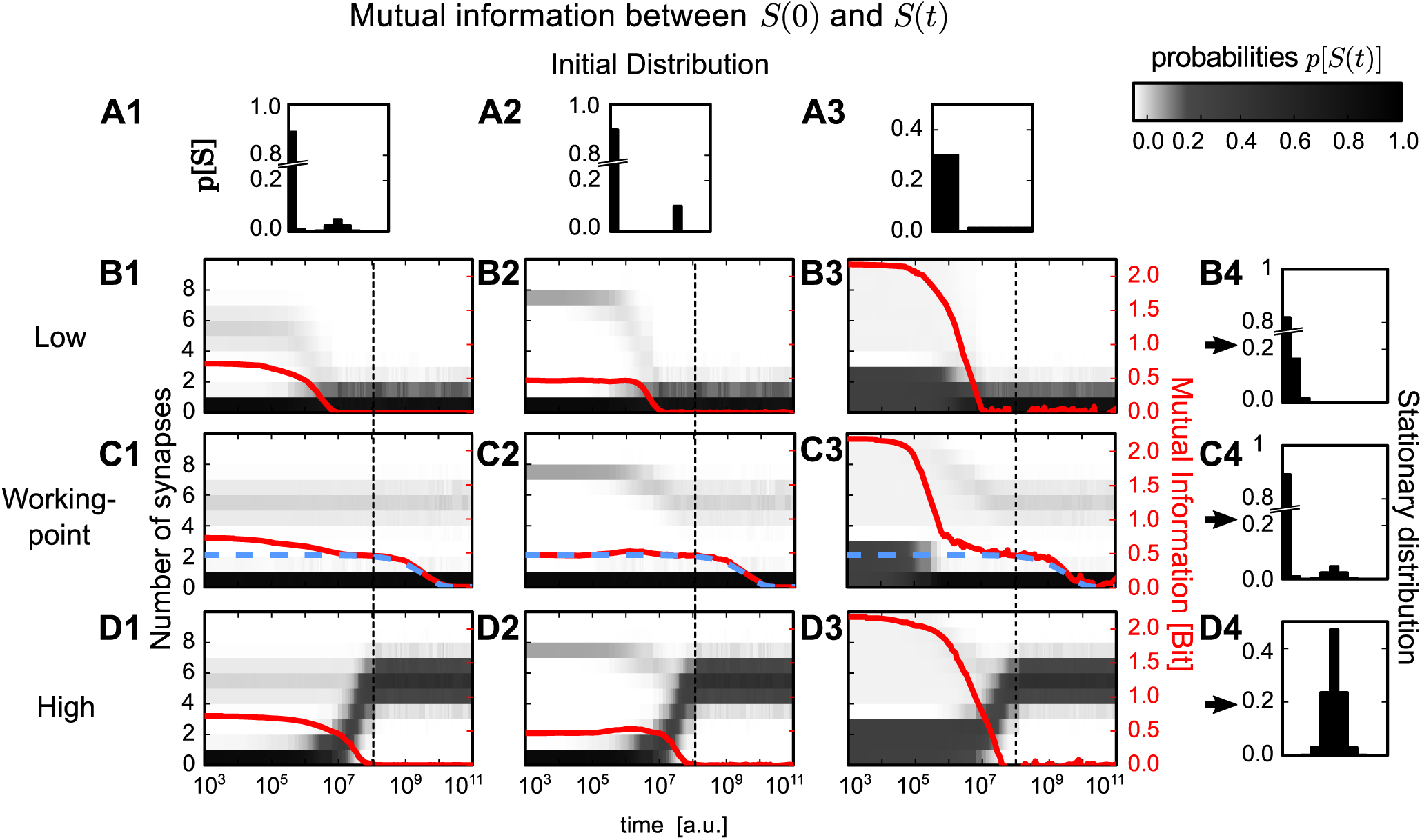
Development of mutual information given different initial and stationary distributions. Three different initial probability distributions (A1–3) and three different stimulation-conditions each yielding a specific stationary distribution (B4, C4 and D4). Dependent on the different combinations of these distributions and conditions, the development of the probability distribution of synapses *p*[*S*(*t*)] differs (gray scale in central panels). This results in different time courses of the mutual information (red) between initial number of synapses *S*(0) and number of synapses *S*(t) at time point *t*. In general, for all conditions, except *wp-*condition, mutual information decays to zero after 10^7^ to 10^8^ time steps. For the *wp-*condition (panels C1–3) the same decay occurs, but to a non-zero plateau. On this plateau information about the initial number of synapses is preserved up to 10^9^ to 10^10^ time steps. The height and duration of this plateau can be reproduced by analytical methods from a simplified system (blue dashed line, see Models and Methods).

Note, although we use a population of connections to evaluate mutual information, this information is stored by each connection on its own as connections have no means of interacting in the model. The use of a population is necessary to estimate the time evolution of the distribution p[S] which describes the number of synapses on a single connection as a random variable. To apply mutual information also the initial condition is treated as a random variable which follows a given probability distribution, although each distribution has only one initial condition and populations with different initial conditions are separately tracked (see Materials and Methods).

Here, we used three different initial conditions for the starting distributions: (1) the *wp-*distribution (Fig. 3 A1); (2) a two peaked distribution with probabilities *p*[*S* = 0] = 1 − *C* and *p*[*S* = 7] = C (Fig. 3 A2); as well as (3) a distribution which is uniform in each peak (Fig. 3 A3) where the summed probabilities are 1 − *C* for the lower peak (*S ∈* 0, 1, 2) and *C* for the upper peak (*S* > 3).

Using these three initial conditions, the three stimulation-conditions from above (see Fig. 1 B) were investigated. Each condition yields after convergence a different stationary distribution (see Models and Methods for more details): (i) *low* induces a single peaked Poisson distribution (Fig. 3 B), (ii) *wp* yields the working-point-distribution (Fig. 3 C), and (iii) *high* leads to a Gaussian distribution (Fig. 3 D).

For *low* and *high* the mutual information and, therefore, the information about the initial distribution, decays after a short period to zero (10^7^ to 10^8^ time steps; Fig. 3 B and D). This period **corresponds to the time scale of the synapse-formation probability rate *b* = 10^−8^/time step, which is** the intrinsic time scale of a compound-connection. Thus, information cannot be stored longer than the average duration of a single synapse turnover. For the *wp-*condition, this decay also occurs after the same period (Fig. 3 C). However, in contrast to all other cases, information does not decay to zero but converges for a longer time to a constant non-zero value. After another long period, mutual information starts again to decay until it finally reaches zero and all information about the initial number of synapses is ‘forgotten’. The period between the first and the second decay provides a plateau indicative of prolonged information storage which will be further investigated next.

### The collective dynamics of multiple synapses encode long-lasting information

The intriguing effect that systems, which converge to the *wp-*stationary distribution, can retain a longer lasting memory of the initial conditions across two phases, while others loose it after a first, early decay can be intuitively understood as explained in the following (Fig. 4 A-C), but this will later be expanded by a mathematical argument using a two state model, too.

**Figure 4.**
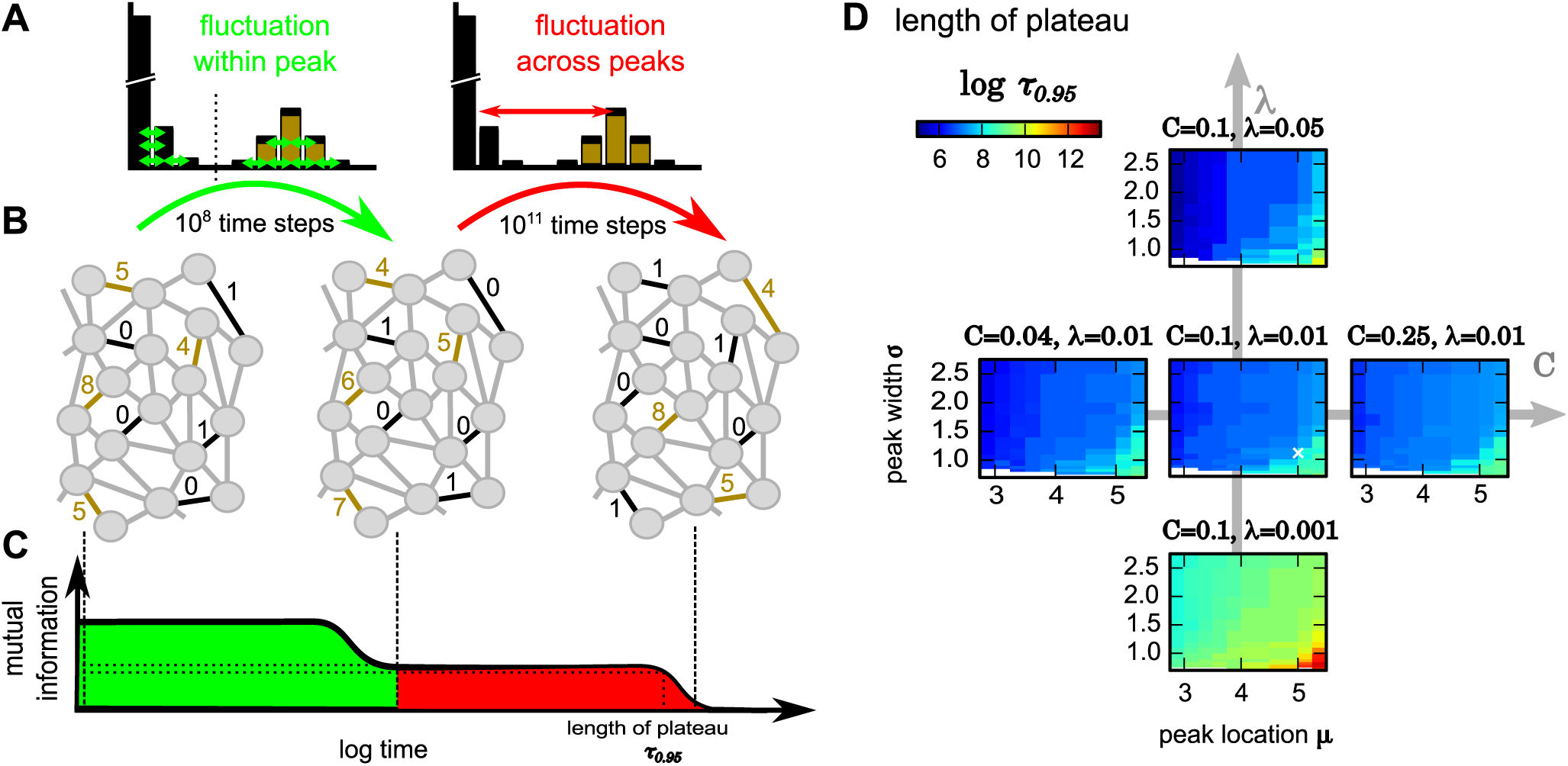
Schematic diagram of the dynamics underlying long-lasting memory and influences of parameters on storage time. (A-C) The different phases (green, red) of decay in mutual information can be explained by *within peaks* versus *across peaks* redistribution of synapses. (D) The period until the plateau of mutual information has decayed to 95% of its initial value (color code, on a logarithmic scale) depends on the parameters of the bimodal *wp* stationary distribution of synapses. White cross in center panel marks parameters used in Figures 2 and 3.

For the bimodal *wp-*distribution, but not for the other ones, the population of compound-connections undergoes two distinct phases of information loss (green and red in Fig. 4 A-C): First, every compound-connection retains its basic characteristic and fluctuates around one of its two stable fixed points at *S* = 0 or at *S* = *μ*. A connection with, for instance, five synapses (*S* = 5; Fig. 4 B, top left) may change to one that has four synapses (*S* = 4; Fig. 4 B, middle) but will not loose all synapses as it experiences a stronger drive towards the upper fixed point *μ* = 5 (brown compound-connections). Likewise a weak connection with one synapse only may drop to zero but not move up to a high number of realized synapses (black). Thus, the basic connection pattern stays the same (compare brown and black compound-connections in left and middle panels in Fig. 4 B). Synapse numbers are redistributed *inside* each peak of the bimodal *wp-*distribution (Figure 4 A, left; first phase). Hence, the mutual information between initial condition and this stage decays but remains above zero: Connections “remember their identity” and a plateau is formed (Fig. 4 C).

As the connections experience a drive towards their current stable fixed point, they approach the basin of attraction of the other stable fixed point only very rarely. This leads to very few transitions from one peak to the other. Thus, changes of the peak can only be expected at much longer time scales. At these time scales, the peak identity is lost and a compound-connection can take any value of realized synapses regardless of how it had been originally initialized. Synapse numbers are finally redistributed *between* peaks (Figure 4 A,B right; second phase) and mutual information drops to zero.

Panels C1 and C2 in Figure 3 represent two interesting cases supporting this view. In C2 the initial condition had been a distribution with two solitary peaks. Hence with respect to the peak *position* such a system is fully determined right from the beginning. Thus, as long as peak identity information is retained there cannot be any drop in mutual information, which is the case until the late and only drop. Until then the system had indeed retained the information about peak positions. Panel C1 started with the *wp-*distribution; the peaks of which have a certain width. Thus, there are indeed various choices for redistributing the synapses *within* each peak and, as a consequence, there is a first phase with a slow drop in mutual information happening until the final information decay, which again corresponds to a mixing of synapses *between* peaks.

### Approximation by a two state model reveals determinants for time scale

To test whether the long-lasting information indeed corresponds to information about the peaks of the bimodal distribution, we designed a reduced two-state model which only describes the transitions between the two peaks. This reduced system can be analyzed by using analytical methods (see Models and Methods for details). We matched the parameters of the reduced model to the initial and stationary distributions of the full system and compared the development of their mutual information. To set the time scale correctly, we searched for the two states with the least probability flow between the two peaks of the *wp-*distribution. The lower one of these two states is the minimum of the *wp-*distribution 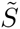. Matching the probability flow of these two states with the probability flow in the two state model yields a transition rate 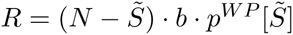 (see Eq. 8), which we use for our simulations.

Indeed, the two-state model reproduces with high accuracy the second decay of mutual information for all three initial conditions (blue dashed line in Fig. 3 C1–3). Therefore, the information, which is stored for long periods, has to be the information about the “identity” of each compound-connection about the peak it had been initialized in. The information about the specific initial number of synapses is lost during the first decay.

These results imply that the long-term storage in the model is not accomplished by single persistent synapses, but rather by the collective dynamics of all synapses of a compound-connection. Remarkably, single synapses can be created and deleted without perturbing this type of storage.

The transition rate *R* of the two state model also provides insights into the typical time scale for memory storage in these systems. Interestingly, this rate only depends on the minimum probability between the two peaks of the stationary *wp-*distribution 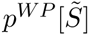 This probability is influenced by the different distribution parameters. Accordingly, storage time is longer for sharper and stronger separated peaks of the final distribution of synapses. Intuitively, with sharp, widely spaced peaks there are less ‘chances’ for a given compound-connection to be at a state, where it can make a ‘jump’ to the other peak, as compared to a situation when the peaks are wider and/or closer together.

We evaluated this by assessing the time scale ro.95 at which the mutual information plateau decayed to 95% of its initial value (after the first decay), while varying all parameters that influence the shape of the stationary distribution: the mean and variance *λ* of the lower peak, the mean *μ* and variance *σ* of the upper peak, and the weighting *C* between the peaks (see Fig. 1 D). All other parameters are the same as for Fig. 3 C1.

The results confirm that in the *wp-*distribution the decay time scales τ_0.95_ gets larger for narrower peaks (small *λ* and σ) and when the upper peak is farther away from the lower one (larger *μ*). The weighting *C* between peaks seems to have a minor influence. This can be also shown by the reduced two-state model as *C* influences the stationary probabilities linearly while all other parameters contribute exponentially (see Models and Methods).

On the other hand, the fact that R only depends on the minimal probability 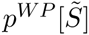 has another interesting consequence: All distributions, which have the same minimum at 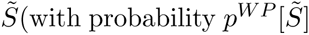 (with probability 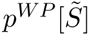 and the same *C*) will always yield at least the same time scale for long-term information storage. Thus, the actual shape of the peaks is unimportant, as long as they give rise to the same minimum in between them (see Supporting Text S1).

Therefore, we can conclude that collective dynamics of all synapses in a compound-connection, which gives rise to bimodal distributions of the number of synapses, also stores information longer than expected from single synapse dynamics.

### Learning can be faster than forgetting

Up to now, we have analyzed how information decays in a system of compound-connections with stochastic synapses. Hence, we had investigated storage time and information loss. On the other hand, so far, it is unclear how this information can be encoded into the connectivity of such systems. This can be achieved through learning as discussed next.

First, before learning, the system has reached its *wp-*stationary distribution (Fig. 5 A, left) corresponding to the steady state under resting state activity. Then, learning is triggered where a changed stimulation would elicit higher activities at some connections and lower activity at others, e.g. due to increased excitation and inhibition. This is modeled by applying either *high-* or *low*-stimulation, or – as a control case – the system just stays in the *wp-*condition (Fig. 5 A, center). Here, the learning phase is always long enough such that the new stationary distribution is reached (Fig. 5 B, green color code). Afterwards, learning stops and the system reverts again to the *wp-*condition (retention phase; Fig. 5 A, right).

**Figure 5.**
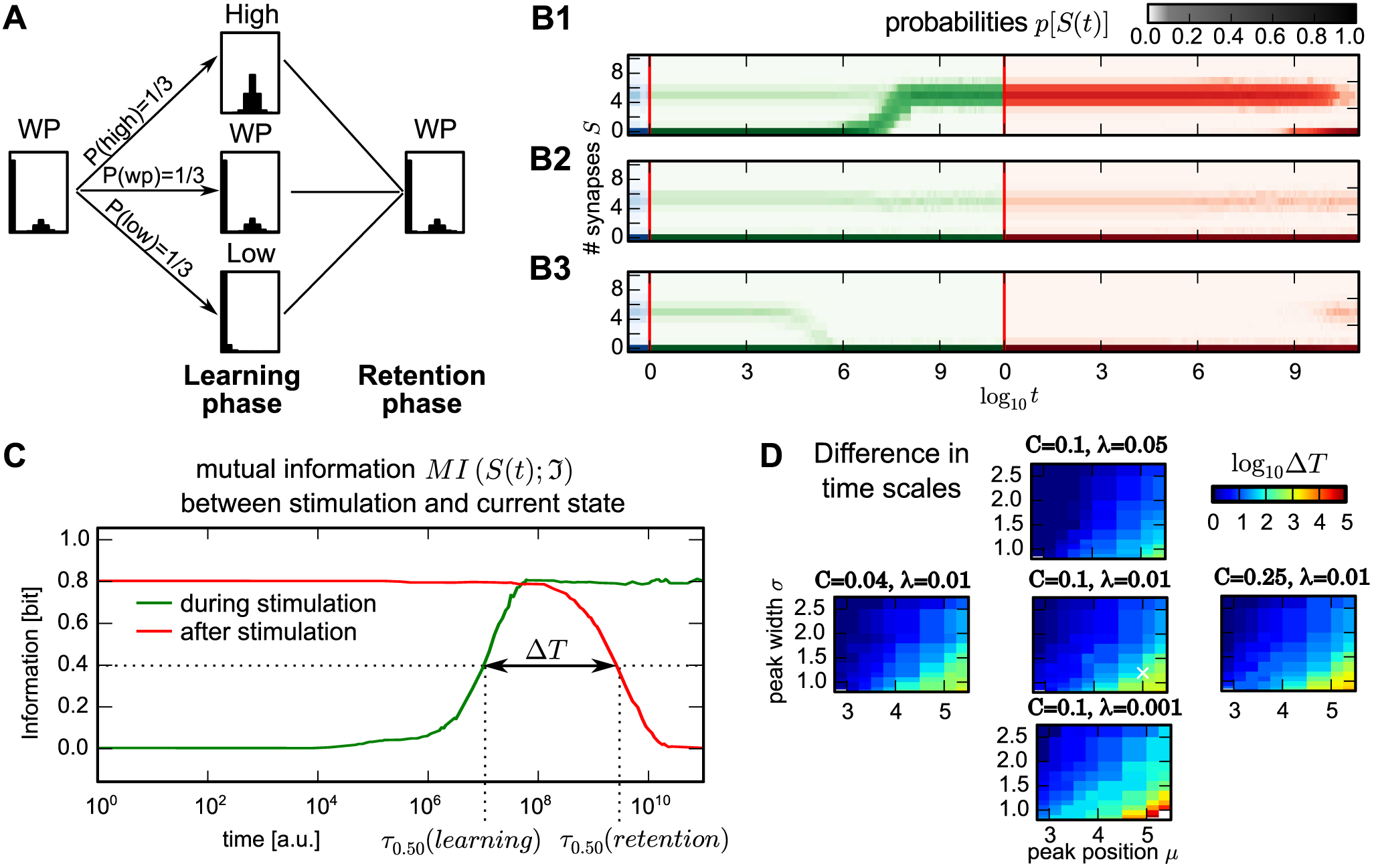
Compound-connections enable learning orders of magnitude faster than forgetting. (A) Paradigm to investigate learning and forgetting: A group of compound-connections is brought from the initial *wp-*condition to either the *high* – or the *low*-condition by applying external stimuli (learning phase). After reaching the corresponding stationary distribution, all conditions are reverted to the *wp-*condition (retention phase) and simulated until the system reaches the stationary *wp-*distribution. The *wp-*only case serves as control. (B1-B3) The time course of the probability distribution *p*[*S*(*t*)] (color code, green: learning phase,; red: retention phase) for the three different cases from (A): B1: *high,* B2: *wp* and B3: *low.* (C) The mutual information between the number of synapses *S*(*t*) and the stimulation-condition during learning phase (green) and retention phase (red) reveals that learning is much faster than forgetting. Dashed lines: time scales τ_0.50_, when the distribution of the number synapses shares 50% of the information of the stimulation-condition. (D) Influence of the distribution parameters on the difference between time scales of learning and forgetting *ΔΤ* measured by the ratio between τ_0.50_ (color coded, logarithmic scale). White cross in center panel marks parameters used in other panels and Figures 2 and 3.

The two different “learning” stimulations (low, high) induce development of the probability distribution *p*[*S*(*t*)] to their characteristic distribution (green phase in Fig. 5 B1,B3), while for the *wp-*condition – as expected – the *wp-*distribution remains roughly constant (Fig. 5 B2). Remarkably, in the *high-* or *low*-stimulation case the system reaches the stationary distribution during the learning phase about 2–3 orders of magnitude earlier than during the retention phase (red phase). We quantify this effect by assessing how much information the number of synapses *S*(*t*) at time *t* shares with the stimulation case (*low, high,* or *wp*) applied to the system.

Thus, we performed 1500 random draws of these three different cases from a uniform distribution with *P*(*low)* = *P*(*wp)* = *P*(*high)* = 1/3, where each drawn stimulation case was used to drive one compound-connection. By this, the stimulation-conditions can be considered as a random variable and the mutual information between *S*(*t*) and the stimulus-condition can be calculated. Note, the actual choice of the distribution for P(*low*), P(*wp*) and P(*high)* does not matter and results remain qualitatively the same. During learning, mutual information grows until the number of synapses *S*(*t)* represents with a high confidence the stimulation-condition applied to the system (green line in Fig. 5 C). By comparing this learning curve with the retention curve (red line), we see that the learning curve has reached its maximum before the retention curve starts decaying. In other words, information is indeed preserved several orders of magnitude longer than the duration the system needs to store it.

Similar to the decay time scale (Fig. 4 C) the difference in time scales between learning and forgetting should also depend on the shape of the finally reached stationary distribution. To quantify this, we determined the time scales τ_0.50_ at which the mutual information between the number of synapses and the stimulation-conditions reaches 50% of its maximal value in the learning and in the retention phase. The difference between these time scales ΔΤ can be determined by the ratio between τ_0_._50_ (*retention*) and τ_0.50_(*learning*) (Fig. 5 D). As observed for the decay time scale τ_0.50_ (Fig. 4 C), the ratio increases if the two peaks of the bimodal stationary distribution are narrower (small *λ* and *σ*) and further apart (larger *μ*). Also in this case the weighting *C* between the peaks has only a minor influence.

The difference in time scales of learning and forgetting can even be observed for learning with single peaked distributions which are piecewise uniform, i.e. for the broadest or sharpest sharpest possible peaks (see Supporting Text S1). This again highlights that the presented results generalise to a much broader class of distributions.

In conclusion, the interactions between multiple synapses in a compound-connection lead to a fast encoding of information, which is then preserved for longer periods. In the final step we will compare these dynamics with data from *in-vivo* experiments [11].

### Synapse-dynamics in compound-connections is consistent with long-term dynamics of dendritic spines

A similar difference of time scales between learning and forgetting is also observed in experiments investigating the dynamics of dendritic spines; the morphological correlates of excitatory synapses in cortex. For example a few days of training, sensory enrichment, or deprivation can influence the spine turnover several months later [11, 12, 13]. As the dynamics of the synapses in the model are comparable to spine dynamics in experiments, next we will reproduce the experimental data from the motor learning tasks of Xu and co-workers [11].

In these experiments, animals were selectively trained in an early (postnatal month one) and/or a late motor training phase (month four). Some animals were only trained during the late training phase (late-only paradigm; Fig. 6 B), some were trained with the same task during early and late training phase (retraining paradigm; Fig. 6 C) and some did not receive any training at all (control paradigm; Fig. 6 A). For all groups the number of created and removed spines had been tracked. When comparing changes during the early training phase, motor training increases spine creation as well as removal compared to the control paradigm (Fig. 6 D, red bars). During the late training phase (Fig. 6 E) a similar increase in spine turnover emerges in the late-only paradigm (compare red bars in Fig. 6 E right panels with Fig. 6 D right panels). However, animals in the retraining paradigm, which had already experienced training before, show in this case spine creation and removal at control levels (compare red bars in Fig. 6 E left with center panels), which indicates that the early training has left a long-lasting trace in the connectivity [11, 12, 13].

**Figure 6.**
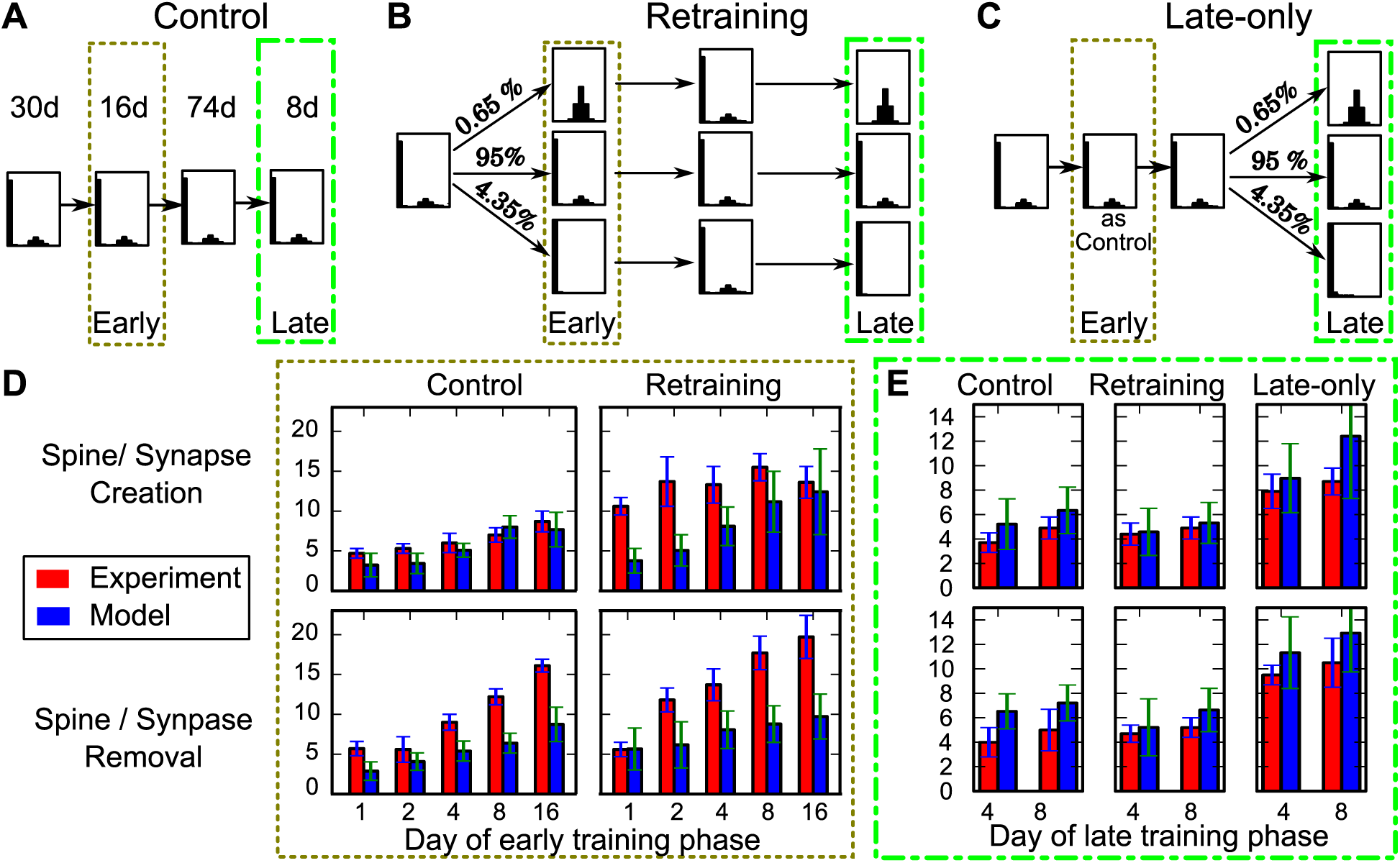
The model reflects long-term effects of spine dynamics. (A-C) Paradigms to test the influence of an early training phase on the spine/synapse dynamics in a late training phase. Durations of the respective phase for all 3 paradigms are indicated above (same for all paradigms). (A) Control paradigm where the system remains in the *wp-*condition (receiving no learning stimulus). (B) Retraining paradigm: Starting with the *wp-*condition, then change stimulation in the *early* phase dividing the population into 4.35% receiving *low*-stimulation and 0.65% of compound-connections receiving the *high*-stimulation. The rest remains unaffected and stays in the *wp-*condition. Afterwards stimulus is withdrawn and the deletion probabilities of all compound-connections are set to render the *wp-*distribution (similar to Fig. 5). After another 74 days a second (*late*) phase with the same stimuli to the same connections as during the early phase is applied. (C) Late-only paradigm: Starting with the *wp-*condition only in the *late* phase *low-* and *high*-stimulations are provided. Note, the middle pathways in (B) and (C) are identical to the control condition (A). (D) Comparison of spine creation (upper panel) and removal (lower panel) of the *early training* phase in experiment (red bars [11]) with the corresponding values from the model (blue bars, mean ± SEM). In this phase the late-only paradigm is identical to the control condition. (E) Creation and removal of synapses in the late training phase are comparable for the control and the retraining paradigm. However, without early training an increased turnover is observed. This corresponds exactly to the dynamics of dendritic spines observed in experiment and indicates the long-lasting trace which the early training leaves in the connections. Here, *N* = 5 and *σ* = 1.0.

In our model these experimental paradigms can be reproduced. For this, again, we take the *wp-*condition as the basic distribution and emulate the process of motor learning by exposing very small sub-populations of compound-connections to *high-* or *low*-stimulation-conditions.

For the *control case,* connections are left in the *wp-*condition during the whole experiment (Fig. 6 A). This case was used to adjust the time scale of the model. For this, we considered the complete temporal characteristic of the experimental control case for spine creation during early training phase (red bars in Fig. 6 D, top-left) and compared it to the model dynamics to match the time scales (see Models and Methods for details).

For the *retraining paradigm* (Fig. 6 B) we start the system in the *wp-*condition for 30 days. Then, for the next 16 days, we define the *early phase* by dividing the population into 4.35% of compound-connections receiving *low-* and 0.65% receiving high-stimulation. The rest does not receive a learning signal and stays in the *wp-*condition. After this, the learning signal is withdrawn and the system and all connections are in the *wp-*condition for another 74 days (similar to Fig. 5 A). Finally, we perform a second (*late*) phase of learning for 8 days with the same changes applied to the same sub-populations as during the early phase. Note, the rather small changes of only 5% of the compound-connections applied here correspond to the sparse changes observed in experiments [11, 12].

For the *late-only paradigm* (Fig. 6 C) we maintain the *wp-*condition for 120 days and only then perform, for 8 days, the same subdivision of the population as above (4.35% *low,* 95% *wp,* and 0.65% *high*).

From these simulations, we determined the realized synapses before and during training phases and calculated the number of created and removed synapses for each of the three different paradigms.

The most interesting case concerns the late phase (Fig. 6 E), where the retraining paradigm had experimentally led to only a control-level rate for creating and removing synapses, indicating that a long-lasting trace might exist in the connectivity. If we compare the numbers of created and removed synapses during the late training phase in our model (blue bars, Fig. 6 E) with the experiments, we find that the model reproduces the data almost perfectly. As in the experiments, the synapse turnover during the late training phase in the retraining paradigm is comparable to the control levels, whereas the turnover is increased, when there is no early training. This indicates that also in our model early training leaves a structural trace in the connections (like in Fig. 5), which persists until the late training.

During early training (Fig. 6 D), model and experiment also match but experimental results show larger creation and removal rates. This is due to the fact that animals show an over-expression of dendritic spines during their early life [32, 9] and accordingly also a stronger removal due to homoeostatic processes. This leads to a non-stationary trend in early development, which is not covered by the model in its current state. To explain the early training phase data completely, we would have to combine the here presented approach with homoeostatic structural plasticity [33, 34], which is beyond the scope of this study.

Thus, apart from such non-stationary effects, our model provides a simple explanation for long-lasting traces in the connectivity. Remarkably, these traces do not rely on the persistence of single synapses (different from [12]) or on different time scales of plastic changes during learning and forgetting but are generated by the natural dynamics of such compound-connection systems.

## Discussion

Here we have demonstrated that storing information on an permanently changing substrate, like dendritic spines, can be achieved by the collective dynamics of spines (or synapses). If the dynamics renders the experimentally observed bimodal connectivity distribution, prolonged information storage about the state of the connections in the network is possible. Other (unimodal) stationary distributions do not exhibit this information storage property. Our results instead suggest that transitions to unimodal distributions might happen during learning and will lead to the desired property that learning is much faster than forgetting in such systems. Importantly these dynamics are consistent with the long-term dynamics of dendritic spines in memory related experiments as shown at the end.

The problem of storing information on a variable substrates is inherent to all neural structures. For example, proteins and molecules, which form the substrate for storing information at any single synapse, are sometimes completely exchanged in rather short times, while the synapse’s long-lasting properties persist [35, 36]. Similar to the here presented approach, this is resolved by the collective dynamics of the involved compounds, i.e., the dynamics of their concentrations and reaction rates. Long-term information storage is achieved if they show a bistable or hysteretic characteristic [24, 25, 26, 27, 28, 29]. Thus, such systems do not rely on the existence of any single protein, analogous to the synapses in the compound-connections in our model.

Such a dynamic is, however, only possible when the single entities are homogeneous, i.e. have the same properties. For multiple synapses between two neurons this might not be the case. Here, the location of synapses on the dendritic tree might influence their impact on the generation of action potentials due to dendritic processing [37] or even change their plastic behaviour [38]. In this case, it would not only matter how many synapses are realised but also which ones. Interestingly, recent experiments suggest that the spine volumes, which are related to synapse stability [39, 40], are very similar for synapses between the same axon and dendrite [41], supporting the homogeneity assumption of our model. However, in general, we expect that qualitatively the here shown prolongation of storage time by the collective dynamics of multiple synapses, is unaffected by such synapse differences, as long as they also implement the experimentally measured bimodal distributions.

Prolongation of storage time is, however, accompanied by a decrease in the amount of stored information. Previous estimates of the information, which can be encoded by the existence of a single synapses, predict it to be around 2–3 bits [3]. In our model, this information dissolves during the first decay phase (see Fig. 3). The maximum information, which can be stored for longer periods by the collective dynamics of all potential synaptic locations at one connection is here 1 bit – connected or unconnected. This value further decreases, when these two states are not equally probable.

However, encoding of information by the collective dynamics of the multiple synapses of compound-connections rescues information from fluctuations of the number of synapses. This has at least three advantages: First, information storage does not rely on the persistence of single synapses. Memories are, therefore, robust against synapse loss. Second, the lifetimes of memories will be much longer than estimated from the lifetime of a synapse (as, e.g., in [12, 13]). Third, connections can keep the same degree of plasticity (by the formation probability rate b) during learning and storage phase. This enables the system to learn very quickly when a learning signal is applied.

The latter seems to contradict the suggestion that there are distinct *learning-* and *memory-spines* [12, 40]. However, in the here proposed model, synaptic plasticity leads to very different dynamics for synapses in the lower as compared to the upper peak. Synapses of compound-connections in the lower peak will repeatedly appear and disappear, thus, exhibiting small synaptic weights. It is known that small weights are susceptible to large changes induced by synaptic plasticity [42]. As a consequence, the present model inherently contains a population of synapses which resembles *learning spines.* On the other hand, synapses of compound-connections in the upper peak create a rather stable connection relating to *memory spines*. This yields a second population of spines with long life times, which have been proposed to encode long-lasting memory traces. This picture is supported by the fact that the long-term dynamics of dendritic spines observed in experiments indeed match the dynamics of the here proposed compound-connections (Fig. 6). Only at short time scales model and experiment correspond less well and the time course of spine creation and removal during training is faster in model than in experiments. This might be due to an increased formation rate of new filopodia or spines during periods of high activity [43, 44], because this leads to an *activity-dependent* synapse-formation rate. In our study, we had restricted ourselves to a constant formation rate to demonstrate the effect of the bimodal *wp-*distribution on the time scale of information storage without having to account for interfering non-stationary effects at the same time scale. Analysis of such coupled systems, in possible future work, is very difficult and needs to build on the here presented results. Strikingly, as all time scales reported here are proportional to the formation probability *b*, an increase of *b* during the stimulation or learning phase would yield to an even faster learning and a larger difference in the time scales of learning and forgetting.

In summary, the here presented model is to our knowledge the first that shows how information can be acquired and stored on the variable substrate of continuously changing spines (or synapses). To achieve this, encoding exploits the existence of multiple synapses between two neurons and their mutual interactions. This principle may lend itself helping to understand the dynamics of long-lasting memories.

## Models and Methods

### Stochastic model of structural plasticity

In this study we model the connection between two neurons by a number of potential synapses *N* and the number of realized synapses *S*. To mimic the overturn dynamics of dendritic spines observed in experiments, synapses are created and removed randomly. Hereby, new synapses are formed with a constant probability *b* at each of the potential synapses which are not realized and existing synapses are removed with a probability *d^ℑ^*(*S*) which depends on the number of synapses and the current stimulation (see Fig. 1A).

### Stimulation conditions

The number of synapses is, thus, a random variable, which follows the dynamics of a Markov chain with the different numbers of synapses as states. The time development of the distribution of the number of synapses *p*[*S*] is then characterised by the transition matrix between the states which is given by *b* and *d^ℑ^*(*S*). As long as all *d^ℑ^*(*S*) > 0, this matrix is non-negative implying that the Markov chain is irreducible. As connections can also stay in the same state, the Markov chain is also aperiodic. For a Markov chain with these properties, the Frobenius-Perron theorem guarantees that there is a stationary probability distribution *p*[*S*], to which an ensemble of such connections will converge [22, 23].

Our previous investigations of the interaction of activity and synaptic plasticity with a similar structural plasticity rule [23] showed that this stationary distribution can be controlled by applying specific stimulations to the pre- or postsynaptic neuron.

In this study, we simplify these stimulations to three different stimulation conditions: a high, a low and a working-point condition. For each of those conditions, we choose a stationary distribution *p^ℑ^*[*S*] with ℑ ∈ {*high, low, wp*} inspired by our previous results (see Fig. 1B): The working-point distribution will take a bimodal shape like the distributions of the number of synapses which were experimentally found in cortex [20, 16, 19, 18, 17]. The high and the low stimulation stationary distributions follow the shape of the upper and the lower peak of these distributions respectively:

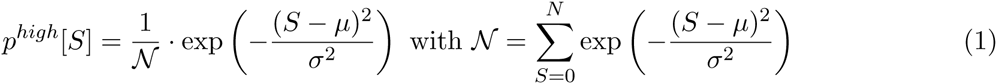

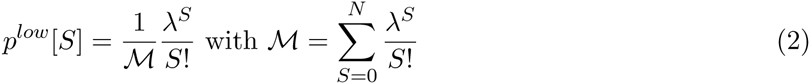

with *μ* and *σ* as the location and width of the upper peak and *λ* adjusting the steepness of the decay in the lower peak. The *working-point* distribution is then given by a mixture:

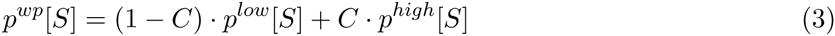

weighted by the connection probability *C* (see Fig. 1D).

### Calculation of deletion probabilities

To construct connections which have the given stationary distributions, we have to determine suitable deletion probabilities *d^ℑ^*(*S*). For this, we use an approximation of the stationary distribution in the described system: When we only allow for creating or removing one synapse at a time ( first step approximation [22, 23]), the transition matrix of the system is

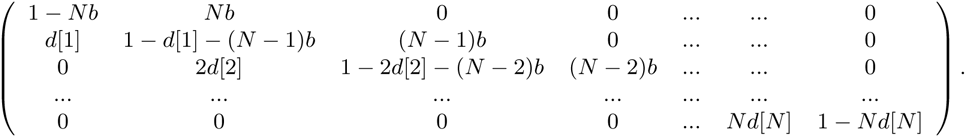

In equilibrium, the probability flow between states *S* – 1 and *S* must be equal (detailed balance) and the stationary distribution *p^ℑ^*[S] fulfils

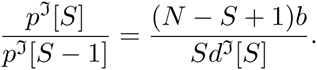

Thus, we choose the deletion probabilities as

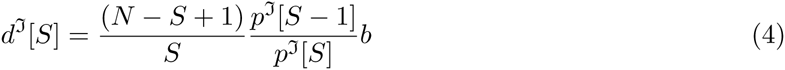

for each stimulation condition. As, by construction, *p^ℑ^*[*S*] > 0, this also yields *d^ℑ^*[*S*] > 0. As described above, this ensures the convergence to the stationary distributions from any initial condition.

### Simulations

**Implementation** The stochastic changes of synapses are implemented in an event-based fashion. Hereby, the time interval until the next removal or creation event for each potential and existing synapse follows a Poisson distribution with different success rates. The number of time steps until the next event can be calculated as the value at which the cumulative probability density function of this Poisson-distribution reaches a random number between 0 and 1. After each change in the number of synapses, all intervals of a connection are recalculated as the success rates for the Poisson distributions change (because of *d*[*S*]). The simulation then proceeds to the next event in the population of connections. Therefore, the calculation time depends on the number of changes in *S* and the time scale can be freely adjusted without increasing simulation times.

To investigate the dynamics ensembles of these connections, we conduct the following simulations:

**Decay of initial conditions – Constant stimulation condition (Fig. 2 and 3)** In these simulations, 500 connections are initialized at the same number of synapses and simulated with the same stationary distribution. This is repeated for all initial conditions. To cover all different time scales of the dynamics, the relative frequencies for all numbers of synapses are saved for logarithmically increasing time intervals.

**Learning scenario – Changing stimulation condition once (Fig. 5)** This paradigm has three phases: (1) In an initialization phase all connections are initialized with the same number of synapses and with the working-point stationary distribution. The length of this phase is long enough that the ensemble reaches the stationary distribution and the initial conditions have no influence any more. (2) In the learning phase, the stationary distribution is switched to either *high, low* or *working-point* condition by adjusting the deletion probabilities. Hereby, each stimulation is selected with a probability *P*(ℑ). Again, this phase is long enough that the stationary distributions are reached. (3) Finally, in a retention phase, the deletion probabilities are switched back to working-point condition again and the population is simulated until it reaches equilibrium. In all three phases the relative frequency of each number of synapses is saved in logarithmically increasing time intervals from the beginning of the phase. The simulation is then repeated for all initial conditions and all possible stimulations.

**Dendritic spine experiments – Changing stimulation condition twice (Fig. 6)** To investigate, whether the data from dendritic spine experiments can be reproduced by the here proposed model, we adapted the stimulation scheme from the *in vivo* experiments [11]:

A population of connection is initialized with the working-point stationary distribution and simulated for 30 days. Then, for the early training and retraining data, the connections are set to different stimulation conditions with the respective probabilities *P*(*ℑ*) for 16 days, whereas for control or naive training data, the stationary distribution remains the same, Afterwards, all connections are set back to working-point conditions and simulated for another 74 days. Then, for the late training phase, the connections are set to the stimulation conditions again. Hereby, the same stimulation as in the early training phase is used for the retraining data.

During stimulation, the occupancy of each potential synapse is tracked, such that we can evaluate the number of persistent synapses or spines from the overlap of occupied locations at at two time points. Comparing this number to the number of synapses at the first and second time point results in the number of created and removed synapses.

**Matching the time scale to experiments** As a first step to compare the results of model and experiment, the number of time steps *T* in the model which correspond to one day in experiment has to be set. To analytically estimate this number we use the fact that, in experiments, under control conditions, one observes about 5% newly formed synapses as compared to the pre-existing ones one day before. Therefore, we estimate the proportion of newly formed synapses under *wp-*conditions depending on T and determine the value of T where it reaches 5%.

For this, it is not enough to just estimate the number of synapses which has been formed, but also the persistence until their next measurement has to be taken into account. To approximate this, we assume that a connection forms at most one synapse during the time interval [0, *T*]. Then, the probability that a synapse is formed at time *t ∈* [0, *T*] and persists until the next measurement at time *T* is given by

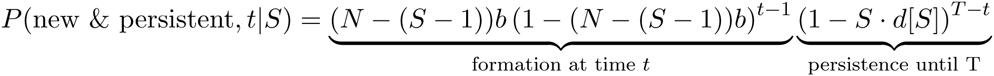

where *S* is the number of synapses after the formation. If we sum this over all possible creation times between 0 and *T*, we get

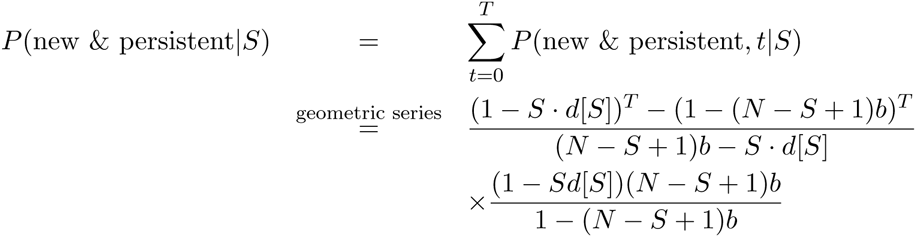

(see Supporting Text S2)

The fraction of newly created (and persistent) synapses is then given by the expected value of this probability divided by the expected value of *S* − 1:

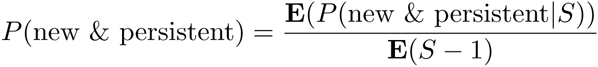

When evaluating this fraction for the used *wp-*distribution (*μ* = 5.0, *σ* = 1.2, *λ* = 0.05, *C* = 0.1, *N* = 5), we find that it first hits the experimental value of 5% at T ≈ 2.3 · 10^7^ time steps (see Supporting Text S2).

We re-evaluated this result by a simulation of the turnover dynamics of populations of our model synapses between day 30 and 46. We compared the proportions of newly formed synapses from these simulations with the values of the control group in experiment and determined the quadratic error. This was repeated for *T ∈* [5 · 10^6^, 30 · 10^6^] in steps of 10^6^ time steps. We found that there is indeed a minimum of the squared error around *T* = 23 · 10^6^ time steps (Supporting Text S2), such that we used this value for our further simulations.

**Determining the stimulation probabilities** Afterwards we simulate a population of connections, each driven by a randomly selected stimulation condition. The size of this population is chosen such that the number of synapses in the initial conditions matches the average number of spines observed in experiment (e.g., for 160 synapses for [11]). These simulations are repeated multiple times (here 8 times which corresponds to the typical number of animals in [11]) to obtain an estimate for the statistical error of the obtained values. These simulations were repeated, while systematically varying the proportion of synapses which receive (*high* or *low*) stimulation, and the fraction of stimulated connections, which received the *high* stimulation (conditioned probability for *high* stimulation).

From these simulations we determine the squared error to the experimental values at the late training phase and the last day of early training, such that the structural traces in model and experiment are based on an equal amount of structural changes. This enables us to determine the best fitting stimulation probabilities *P*(*ℑ*). Interestingly, our best match occurs at only 5% stimulated connections, out of which 13% receive *high* stimulation.

### Reduced two-state system

#### Dynamics and entropy

If the long-lasting information encodes the peak a connection was initialized in, a model of the probability masses in the two peaks should be able to reproduce the second decay of mutual information in the full system (see Fig. 3C1–3). To test this, we use a Markov chain with two states for the lower and the upper peak *p*:= (*p*_0_*, p*_1_) (note this system would also describe the behaviour of a mono-synaptic system). Assume the stationary distribution of this system is (1 − *C, C*). Then, the transition matrix of such a process takes the form:

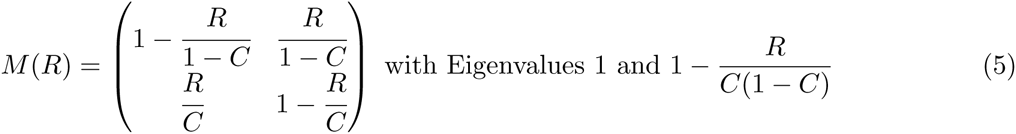

where transition rate *R* adjusts the convergence time to the stationary distribution and is a free parameter. The time evolution of the system with initial condition (1 − *p_init_, p_init_)* can approximately be described as

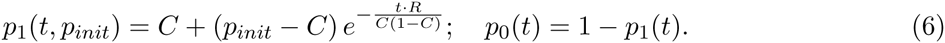

The mutual information between initial conditions and the distribution p(t) evaluates to

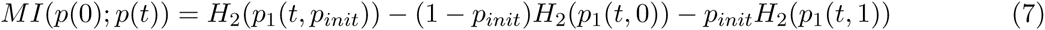

with a two state system entropy

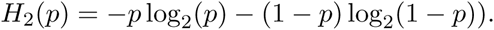

#### Transition rate *R* corresponding to the multisynaptic system

Beside the initial condition and the stationary distribution, the time scale of the two state model has to be matched with the full system. For this we analyse both systems in their steady state:

In equilibrium (i.e., (*p*_0_, *p*_1_) = (1 − *C, C*)) the probability flow between the states of two-state system must be balanced (equal to *R* by construction):

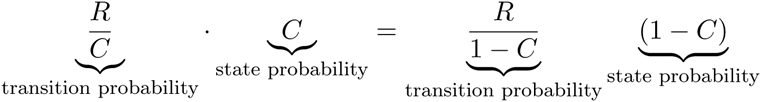

If we again assume that in the multisynaptic system only one synapse is created or removed at a time, the same holds for the probability flow between the upper boundary of the lower peak 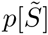 and the lower boundary of the upper peak 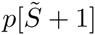 The probability flow between the boundary states is:

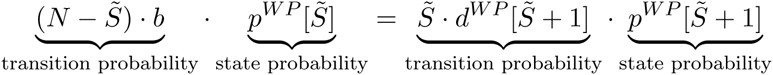

As this is the only probability flow between the two peaks in the multisynaptic system, a corresponding model of the two peaks should have the same probability flow:

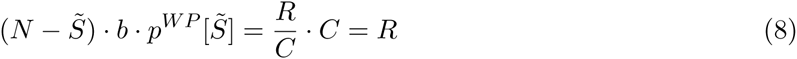

Note, equation 8 can also be used to quantify the advantage of multiple synapses:

Compared to a mono-synaptic system with the same building probability, the information decay is 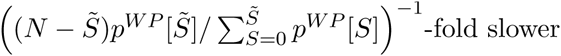-fold slower.

#### Information theoretic measures

To quantify shared information between two random variables and their probability distributions, the information theoretic concept of mutual information can be used. Here, we use it for two different analyses:

**Mutual information between the number of synapses at different times (Fig. 3)** If we regard the number of synapses at time step 0 (i.e., *S*(0)) and at timestep *t* (i.e., *S*(*t*)) as random variables, we can evaluate the mutual information between them as

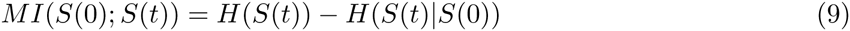

with the entropy 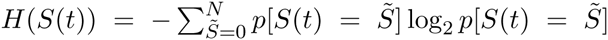 and the conditional entropy 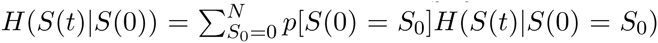 with 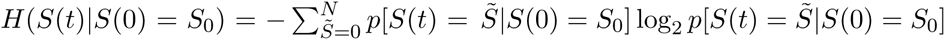

Note, this formulation of mutual information can be evaluated only from simulating the distributions *p*[*S*(*t)|S*(0) = *S*_0_] for each initial condition *S*_0_. The entropy *H*(*S*(*t*)) can then be calculated from the sum of the distributions emerging from different initial conditions *S*_0_ weighted with their probability *p*[*S*(0) = *S*_0_] (i.e., total probability):

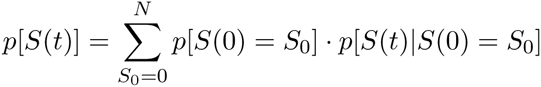

In this way the initial conditions can be chosen freely and the mutual information can be evaluated for different distributions of S(0) from the same simulated distributions *p*[*S*(*t)|S*(0) = *S*_0_].

**Mutual information between number of synapses and stimulus (Fig. 5)** Similarly, the mutual information between the number of synapses at time t and the applied stimulation condition can be evaluated as

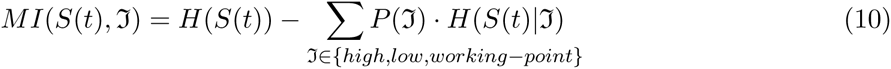

where *P*(*ℑ*) is the probability that a connection experiences a stimulation condition *ℑ ∈* {*high, low, WP*} during the learning phase and *H*(*S*(*t)|ℑ*) the conditional entropy of the distribution *p*[*S*(*t)|ℑ*] when the stimulus is known.

For each stimulation, again each initial condition can be simulated separately. Afterwards the distributions *p*[*S*(*t*)] and *p*[*S*(*t*)|ℑ] are calculated as weighted sums of the simulation results. Here, the initial conditions and *P*(*ℑ*) can be chosen freely, although the initial conditions should have decayed during the initialization phase.

## Supporting Information

### Supporting Text S1

**Learning and forgetting with other stationary distributions** Here we investigate, how the presented results generalize to a broader class of stationary distributions for learning and forgetting.

### Supporting Text S2

**Results from matching model and experimental data**

## Acknowledgments

We thank Prof. M. Tsodyks for helpful comments on earlier versions of the manuscript.

This research was partly supported by the Federal Ministry of Education and Research (BMBF) Germany under grant number 01GQ1005B [CT] and 01GQ1005A [MF, FW], as well as the Göttingen Graduate School for Neurosciences and Molecular Biosciences under DFG grant number GSC 226/2 [MF].

